# Motion and physiological noise effects on amygdala real-time fMRI neurofeedback learning

**DOI:** 10.1101/366138

**Authors:** Lydia Hellrung, Viola Borchardt, Florian N. Götting, Jörg Stadler, Claus Tempelmann, Philippe N. Tobler, Martin Walter, Johan N. van der Meer

## Abstract

Real-time fMRI neurofeedback allows to learn control over activity in a localized brain region. However, with fMRI, physiological factors such as the cardiac cycle and respiration interfere with the measurement of brain activation. In conventional fMRI studies this is usually mitigated by inclusion of motion parameters and/or physiological parameters as nuisance regressors at the analysis stage, allowing to correct for and filter out such confounders. In real-time fMRI, however, such an approach is not routinely feasible due to the necessity to process all signals during the runtime of an experiment. The absence of on-line correction can therefore compromise real-time fMRI study outcomes reporting volitional self-regulation capability as BOLD signal changes. This is especially true for BOLD signal changes in subcortical regions situated close to blood vessels or air cavities, such as the amygdala. We therefore aimed to establish the effects of motion, heart rate, heart rate variability, and respiratory volume on learning effects, which means here an increase in BOLD signal change over NF training, in an amygdala neurofeedback experiment. Specifically, we investigate motion parameters from two emotion regulation studies - performed at 3T and 7T scanners - and additionally acquired physiological variance for the latter one. Our results revealed differences in these parameters between groups and especially between regulation and resting periods within each participant. However, strictly considering these parameters as nuisance regressors in data analysis revealed that the learning of volitional self-regulation of the amygdala is not driven by motion and physiological changes. As validation of our real-time findings, we compare them to the gold standard of assessment of motion and physiology from the Human Connectome Project. Based on this, we recommend to carefully report neurofeedback study results including physiological nuisance regression. To our knowledge, this is the first study investigating the effects of motion and physiological noise correction on neurofeedback BOLD effects.

## 1 Introduction

Real-time fMRI (rt-fMRI) allows to learn volitional self-regulation and has been performed with numerous brain regions (Sitaram et al., 2016). A region of interest (ROI) is determined via either an anatomical or a functional localizer, which is subsequently used to extract the blood oxygenation level dependent (BOLD) signal for neurofeedback (NF). The presence of the neurofeedback signal thereby allows participants to exert control over BOLD activity through training. Unfortunately, BOLD activity changes can also be induced by motion or physiological changes, such as changes in heart rate and/or respiration (Kasper et al., 2017; Parkes et al., 2018). Respiration and heart rate could impact BOLD in two ways – first by pulmonary and respiratory motion interacting with the localized magnetic field distortion causing signal distortion and dropout; second by changing the blood oxygenation levels all over the brain, impacting the BOLD contrast at a physiological level. To clearly separate NF learning effects from such confounds requires to fully consider these confounds during data acquisition and analysis. So far, in conventional fMRI research head motion can be extracted from imaging data and respiration and the pulsation of blood vessels can be acquired in parallel to imaging with appropriate sensors (respiration belt for breathing and electrocardiogram (ECG) or pulse oximeter (PPU) for heart beat). Nowadays, this is a highly-recommended procedure. Taking these data into consideration for analysis is straightforward with the appropriate toolboxes, such as the PhysIO Toolbox (Kasper et al., 2017). These toolboxes extract nuisance parameters from a complete data set. The extracted parameters can then be used for offline fMRI data analysis to regress out motion and physiology-related variance in the BOLD signal.

For rt-fMRI it is more challenging to properly consider such nuisance variables due to the component of the *on*-*line* analysis, with (by necessity) reduced amount of available data points during the runtime of an experiment. The effects of motion in on-line analysis are usually considered in terms of an on-line realignment of the image data. Such approaches are either already provided by the sequence reconstruction from the MR scanner (e.g. Siemens or Philips prospective motion correction for EPI imaging) or by on-line preprocessing within the software processing pipeline (Goebel, 2012; Hellrung et al., 2015; Koush et al., 2017a). It is challenging to include rotation and translation parameters in rt-fMRI in analogy to conventional fMRI. However, some software toolboxes allow to integrate motion regressors into the on-line statistics by using the first measured volume as a reference (Hellrung et al., 2015; Hinds et al., 2011; Koush et al., 2017b). In contrast, real-time correction of physiological confounders is currently not possible and it still is a largely open question how they affect neurofeedback results.

Therefore, in this study, we investigate the effects of motion, heart rate variability and respiration on the NF learning effects in rt-fMRI regulation experiments. The amygdala is a ROI of high interest for rt-fMRI neurofeedback experiments. Its capacity for being modulated by means of neurofeedback has been investigated so far in numerous studies including two own previous studies (Brühl et al., 2014; Hellrung et al., 2018; Marxen et al., 2016; Paret et al., 2014; Zotev et al., 2011). Furthermore, amygdala NF training has been proposed as a potential complement to pharmacological interventions for disorders, such as depression (Young et al., 2014) and Posttraumatic Stress Disorder (Gerin et al., 2016). However, the effects of motion, respiration and heart rate could be especially profound for the amygdala due to its close proximity to major blood vessels (Boubela et al., 2015), and its proneness for signal dropout (Deichmann et al., 2002).

Specifically, we need to know how head motion, respiration and heart beat will affect real-time fMRI learning effects, especially for amygdala. The following questions need to be answered:

(1) How do motion and physiological measures (heart rate, heart rate variability and respiration) vary during the neurofeedback experiment, and does they vary as a function of either intervention group (feedback vs. no feedback) or actions within the experiment (rest vs. regulation)?
(2) Do these parameters correlate with learning effects, i.e. increase together with increased capacity of BOLD regulation across several training runs?
(3) And, how does the main outcome of an rt-fMRI study (i.e. the demonstration of BOLD regulation and its transfer) potentially change when motion and physiological parameters have been accounted for?

To answer these questions, we acquired data in a real-time amygdala neurofeedback experiment paying special attention to motion and using respiration belt and pulse oximetry to assess respiration and heart rate, respectively. Moreover, we aimed to test whether the variations within our real-time data fall within the normally expected range from a big sample. To do so, we used motion, respiration and heart rate data from the human connectome project which provides a gold standard for quantifying variability in these parameters. The results of our study speak to the growing NF literature. Importantly, our data clearly indicate that amygdala NF learning effects are not primarily driven by motion and physiological noise, a finding which reinforces the feasibility and reliability of amygdala self-regulation. However, our results also illustrate the necessity to report in detail how rt-fMRI studies control for motion and physiological parameters in the analysis.

## 2 Methods

### 2.1 Participants

At the 7 Tesla site in Magdeburg, 34 participants were recruited to perform either an intermittent NF (7T INT; N=20) or a no feedback version (7T NOF; N=14) task. At the 3 Tesla site in Leipzig, a total of 42 participants were recruited to perform a continuous NF (3T CON; N=18), intermittent NF (3T INT; N=16), or a no feedback version (3T NOF; N=8) task. At both sites, the same experimental NF task has been performed (see 2.3). All procedures were conducted in accordance with the Declaration of Helsinki and were approved by the Ethics Committees of the Faculties of Medicine of the Universities of Magdeburg and Leipzig for the 7T and 3T data acquisition, respectively. Participants gave informed written consent prior to participating in the study.

### 2.2 Experimental procedure MR acquisition

#### MRI data acquisition

Scanning was performed using 12-channel head coil on 3T scanner and a 32-channel head coil on 7T. For functional imaging, the repetition time (TR) was identical (2s) on both scanners to facilitate an identical stimulus presentation. In the 7T experiment, the following parameters were used for functional imaging: echo time (TE) = 20 ms; matrix size =160×160 voxel; bandwidth = 1838 Hz, flip angle = 90°; voxel size = 1.4×1.4×1.8 mm^3^, GRAPPA factor = 3. The used sequence served to corrected for motion and distortion during the reconstruction of the images (In and Speck, 2012). These data have been used for the real-time calculation of the feedback signal. In addition, the sequence provided uncorrected data, which has been used for motion analysis within this study. For 3 Tesla, the following parameters were used for functional imaging: TE = 25 ms; matrix size=64×64; bandwidth = 1953 Hz; flip angle = 90°; voxel size: 3×3×2.6 mm^3^. Here, the real-time motion correction has been applied by our analysis toolbox after data export to the external analysis computer. Prior to the functional imaging in both experiments, T1-weighted images were acquired for anatomical normalization using three-dimensional magnetization-prepared rapid gradient echo (MPRAGE) sequence (sagittal orientation) with selective water excitation and linear phase encoding (Mugler and Brookeman, 1990).

#### Amygdala Region of Interest (ROI) Delineation

The T1 scan was used to demarcate manually the left amygdala. Amygdala masks were drawn individually per participant into T1 data by a neurologist using FSLview^1^ (Jenkinson et al., 2012). The amygdala ROI was co-registered and re-sampled into the participant’s own functional EPI space by performing a short localizer functional scan, co-registering this scan to the participant’s T1 scan, and then applying the inverse transformation on the amygdala mask.

#### Real-time Data transfer and fMRI Neurofeedback Software

The neurofeedback setup at both sites was as previously reported in our study (Hellrung et al., 2018). In short. We were using the in-house toolbox rtExplorer (Hollmann et al., 2011, 2008) and a direct transfer of the data via network connection. The data were sent volume-wise to a network port and stored into the random-access memory of the analysis computer. For 3T, data were motion corrected using the preprocessing module of the in-house software BART (Hellrung et al., 2015), while 7T data were motion corrected during the reconstruction within the MR sequence.

#### Physiological Recordings

At the 7T site, we additionally acquired respiration information with a respiration belt (using a pneumatic respiration transducer from Honeywell 40PC001B1A) and heart-rate information with a NONIN (8600-FO) pulse oximeter on the right index finger. For digital recording and subsequent analysis of physiological data we used an in-house setup, consisting of the hardware "PhysioBox" and the software "Physiolog"^2^. The PhysioBox employs the National Instruments acquisition card USB 6008. The Software "Physiolog" written in Python samples the data at 200 Hz and stores them as CSV file. The acquisition of these data is synchronized with the MR triggers. At 3T site, only motion parameters have been acquired. Our questions upon physiological confounders arose from the experiences we gained from that experiment.

### 2.3 Experimental neurofeedback paradigm

The NF paradigm required participants to generate a positive mood by remembering positive memories during up-regulation (HAPPY) blocks. During counting (COUNT) blocks, participants were requested to count backwards from 100 in steps of 3. During resting (REST) blocks, participants were requested to disengage from the task. All blocks lasted 40 seconds. The three different blocks were prompted by different cues: a red arrow pointing upwards for HAPPY, a blue arrow pointing downwards for COUNT, and a white cross for REST. A thermometer was present during each block and was updated in a different fashion for each experimental group as follows: For the intermittent neurofeedback group (INT), the thermometer remained blank during the HAPPY and COUNT blocks, but was updated at the end for 4 seconds with a value representing the average difference in image intensity between the regulation block (HAPPY or COUNT) and the preceding REST block. For the NOF group, the thermometer remained blank in this time period. Participants of the INT group were instructed to attempt to maximally increase (HAPPY) or maximally decrease (COUNT) the thermometer display. During REST the thermometer remained still at the zero point.

The 40-second blocks of REST, HAPPY and COUNT were repeated 4 times in a single run lasting 8 minutes and 40 seconds (with an additional REST block at the end). In the INT group, this resulted in 8 presentations of the NF information, here, i.e. the current BOLD activity within the amygdala. This information was presented at the end of each regulation block (4 for HAPPY, 4 for COUNT) lasting for 4s, which facilitate NF training. The complete rt-fMRI NF experiment consisted of three such training runs. Preceding the training runs, participants performed a Baseline practice run, which consisted only of four HAPPY runs of 80 s each, interleaved with 40 seconds REST, cued with a red upward arrow and concluded with a 4s presentation of NF information. The Baseline enabled participants to practice up-regulation strategies and get used to the scanner environment. The training runs were followed by a Transfer run, which was identical to the Training runs but did not present any NF information.

Before the experiment commenced, a short 40-second localizer functional scan was performed to enable co-registration of the previously delineated Amygdala Mask into the current functional EPI space.

### 2.4 Validation with HCP data

To gauge our levels of variability in motion and physiology against those normally found during MR acquisition, we compared our data to those of the Human Connectome Project (HCP). We used a mirror of the HCP data stored on Australia’s high-performance computing infrastructure located in Melbourne (MASSIVE) (Goscinski et al., 2014). The HCP data includes cardiac and respiratory signals measured with a pulse oximeter and respiratory bellows (Siemens Physiologic Monitoring Unit). These signals, along with the sync pulse from the scanner, were recorded by the scanner host computer at a sampling rate of 400 Hz.

#### HCP tasks

We chose three different fMRI tasks paradigms from the available datasets: (1) “REST1”, where no explicit task was performed, (2) “EMOTION”, which was a block-design emotional picture task supposed to raise amygdala activity and therefore is relevant for comparison to our paradigm, (3) “MOTOR”, where participants performed tongue or bilateral finger and feet movements. We chose this task as it is likely to provide an upper level of median framewise displacements and physiological variation.

#### HCP data

For median framewise displacement evaluation, data of 822 participants were included that had performed all three tasks. To perform between-group statistical comparisons, participants were subdivided into three groups of 274 each, so that each task comprised disjoint participants’ data. To estimate variation in physiological parameters, a total of 636 participants were selected for whom physiological recording data were available for all three tasks. These were also divided into three groups, resulting in 212 participants per group.

### 2.5 Data processing and parameter extractions

#### 2.5.1 Data preprocessing (fMRI)

Preprocessing of the functional data was performed using SPM12 (Wellcome Department of Imaging Neuroscience, London, UK) and Matlab 2011b^3^. It comprised correction for slice acquisition time within each volume, motion correction (motion correction parameters were extracted and used for further analyses as described below), and co-registration to the T1 scan. Next, DARTEL (Ashburner, 2007) served to normalize functional data to MNI space. Images were resampled with a 1.5 mm3 voxel size, high-pass filtered with 240s filter size, and smoothed with a 6mm kernel. Further, individual amygdala masks were normalized to the standard MNI template using the individual T1-weighted structural images.

#### 2.5.2 Median framewise displacements extraction

To quantify and compare motion between participants, we calculated median framewise displacement (MFD) values, which convert the three rotation and three translation parameters obtained from the SPM realignment step in the preprocessing of the functional data into a single motion value per volume (Jenkinson et al., 2002). Specifically, the voxel-specific head motion is a nonlinear combination of the volume-wise translations and rotations, as well as the voxel's position. Here, the voxel-specific movement is handled via a combination of affine matrices for each rotation and translation, Tt ^−1^. The spatial mean across all voxels is then calculated according to Jenkinson:

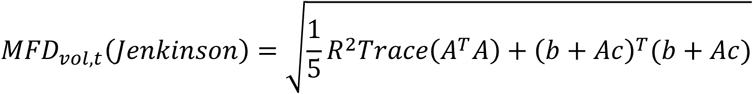

where R=80 mm is assumed as radius of the head, c indicates the coordinates for the center of the volume and A and b are defined as 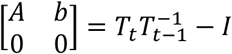 defining rotations and translations (equation and further details in Yan et al., 2013a). The calculation here is based on algorithms from the DPARSF toolbox (Chao-Gan and Yu-Feng, 2010a) using Matlab 2011b.

#### 2.5.3 Physiological parameter extraction

To assess physiology, for all data sets (HCP data files were downloaded for processing), the signals from the physiology station were converted to input text files for the PhysIO toolbox for analysis and modeling of electrophysiological data (Kasper et al., 2017). This toolbox detects breathing cycles from the respiration data and heart beat markers from the physiological data. For this, template models for the pulse complex and respiration are convolved with the acquired signals to detect peaks as heart beats and volume of breath per given time unit.

##### Heart Rate / Heart Rate Variability

We calculated the average heart rate (HR in beats/min) and heart rate variability (HRV) within the different tasks (amygdala regulation for NF data and REST1, MOTOR and EMOTION for HCP data): Furthermore, for our data, we split up these values according to our task conditions (HAPPY, COUNT, REST). The HRV was calculated according to the standard deviation of intervals between the single heartbeats:

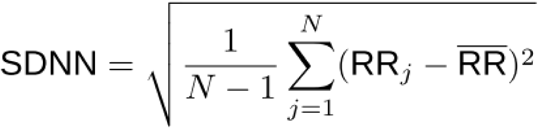

where RR_j_ denotes the value of j’th RR interval and N is the total number of successive intervals.

##### Respiration

To assess respiration, we used the output of the TAPAS toolbox encoding for respiratory volume per unit time (RVT) (Kasper et al., 2017) per scan. We then calculated the average RVT within the different tasks and conditions of our task.

#### 2.5.4 Construction of motion and physiological regressors for GLM

To account for motion and physiology in the analyses of the fMRI data, we constructed regressors that can be used to regress out effects from the functional data. In order to project out motion from the fMRI data, we constructed a matrix consisting of columns containing the realignment parameters, the framewise displacement of the realignment parameters, and scan-nulling regressors. The scan-nulling regressors were constructed by the MFD values and applying a threshold of 0.5 mm to identify functional volumes in which motion exceeded this threshold (Lemieux et al., 2007). A scan-nulling regressor is then constructed for each volume in the interval of −1 to +2 before/after that volume. We used the implementation in the covariate regression step of the DPARSF toolbox to implement the formation of the scan nulling regressors (Chao-Gan and Yu-Feng, 2010b). We chose the named definitions of MFD and scan nulling for this study since it has been shown as reliable and valid method in comparison to other comparable approaches (Yan et al., 2013b)

To account for physiological confounds, we formed a matrix consisting of 8 regressors. Six were RETROICOR regressors (Glover et al., 2000) to model for cardiac and heart beat phase; we used a first order model for cardiac phase, a first order model for respiratory phase, and a first order interaction model. To account for BOLD variations in the imaging data, these three phase models were expanded using a sine and a cosine function as outlined in the RETROICOR methods paper (REF) and the PhysIO toolbox paper (REF), yielding 6 regressors. In addition to the 6 RETROICOR regressors, we included two additional regressors that directly model for HRV and RVT. The HRV regressor was convolved with the cardiac response function (Chang et al., 2009) and the RVT regressor was convolved with the respiration response function (Birn et al., 2006). This is parallel to the approach as outlined in the PhysIO toolbox (Kasper et al., 2017).

#### 2.5.5 Amygdala ROI extractions

To assess whether changes over NF training time in the amygdala regulation capability, i.e. the % signal change in BOLD activity in each training run, could be explained by head motion and/or physiological noise, we constructed three different general linear models (GLM) in SPM12 on single-subject level. All GLMs included eight regressors modelling HAPPY and COUNT conditions for the three training and transfer runs and one regressor modelling HAPPY for the Baseline run. The REST condition was treated as implicit baseline, as participants were instructed to disengage from the task and inclusion of this regressor would have led to a singular design matrix. The GLMs differ in the number of nuisance regressors included in the models: (1) without any nuisance regressors (GLM NO NUISANCE), (2) with six motion regressors and scan-nulling regressors as described in 2.5.4 (GLM + MOCO), and (3) with six motion regressors, scan-nulling regressors and eight physiological nuisance regressors as described in 2.5.4 (GLM + MOCO & PHYSIO). Since physiological data has been acquired for 7T data only, the latter GLM was conducted for 7T dataset only. All single-subject GLMs served to estimate BOLD signal change (% signal change) of HAPPY vs REST for each run (Baseline, Trainings, Transfer) within the individual amygdala ROI. This was calculated by extracting beta parameter estimates from the GLM result files from the HAPPY condition and the implicit baseline regressor as estimates during REST. We averaged the extracted values from the amygdala ROI mask and divided the averages from HAPPY by those from REST.

### 2.6 Statistical analysis

#### Median framewise displacements and physiological parameters

We performed statistical comparisons of MFD values, HR, HRV and RVT between the groups (INT, CON), condition-wise (HAPPY, COUNT, REST), and within-subjects along the runs (Baseline, Run1, Run2, Run3, Transfer) to assess the dynamics of motion. We were using ANOVA functions for between- and within-subject comparisons respectively provided by the Statistics toolbox in Matlab 2016b. For all ANOVA tests revealing significant differences, we performed Bonferroni-corrected post-hoc tests for further insights. To investigate the relationship between motion and amygdala regulation capability, we calculated Spearman correlations between MFD per run with the extracted amygdala % signal change (see 2.5.5) from GLM NO NUISANCE during the respective run using SPSS (IBM SPSS Statistics 25). Calculations of effect sizes, Cohen’s d, have been performed for ANOVAs with multiple groups, based on group means (Jacob Cohen, 1988, p. 273) or for within group comparisons by subtracting the means and dividing by the pooled standard deviations.

#### HCP data validation

To assess group differences in MFD, HR, HRV and RVT within the HCP data between the different tasks, we performed 2-tailed independent sample t-tests on the averaged values from each participant per task using the Matlab 2011b Statistics Toolbox.

#### Interaction of amygdala regulation task with motion and physiological confounds

To compare within-subjects’ differences in amygdala BOLD activity between the three GLMs we transferred the amygdala data for each participant and run into R (R-project R3.4.0). Specifically, we applied a mixed effect model using REML with random slopes and intercepts for each run and each type of GLM. We modelled the main effects of GLM type, run, and their interaction as within-subject factors. These analyses allowed us to investigate whether estimated learning effects from real-time fMRI NF reflect motion and physiological confounds. In addition, we the analyses allowed us to investigate whether statistical group differences in NF learning are affected by noise correction.

## 3 Results

### 3.1 Motion differences (How does motion vary during the neurofeedback experiment?)

To assess differences in head motion we calculated MFD values for all groups from both amygdala NF neurofeedback studies. In detail, we compared following aspects.

#### 3.1.1 Overall group comparisons (Does motion vary as a function of intervention group?)

Our two independent studies revealed differences in median displacement values for head motion between feedback and control groups (3T: ANOVA F(4,8)=10.9, p=.002, Cohen’s d=4.2; 7T: ANOVA F(4,4)=16.24, p=.01, Cohen’s d=6.8). Bonferroni corrected post-hoc t-tests revealed higher motion in all NF groups compared to CON groups (3T: mean INT: .074±.003, CON: .079±.003, NOF: .058±.008, INT vs. NOF and CON vs. NOF p<.001; 7T: mean INT: .079±.002, NOF: .055±.005, INT vs. NOF p<.001). The differences in MFD values are illustrated along the columns in Figure 1 where the black lines indicate the group averages.

**Fig. 1.**
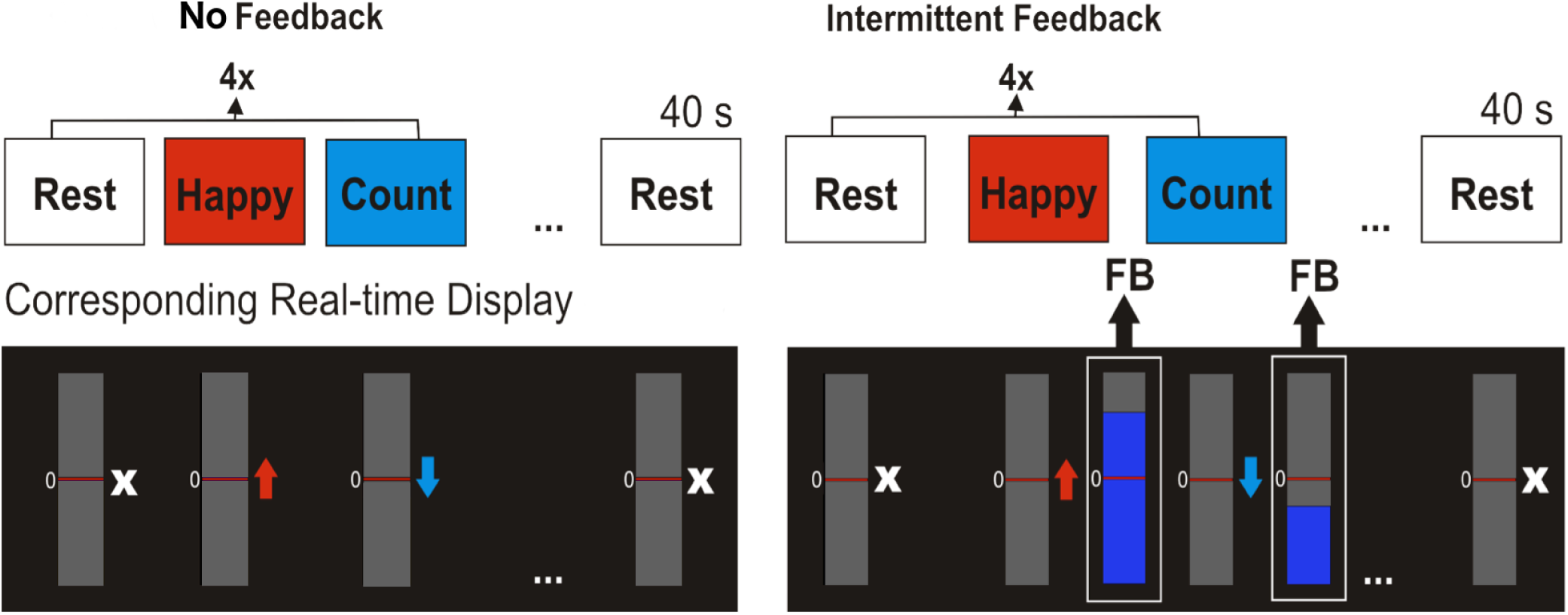
NF paradigm for amygdala self-regulation: Colored arrows or a white cross indicated the current regulation block. The intermittent group (INT) received feedback about their current amygdala activity every 40 s at the end of the regulation block. The control group (NOF) did not receive any feedback throughout the experiment. Overall, the participants repeated this paradigm in three training runs and one transfer run without any feedback.

Given that the central moment tested above revealed differences between the groups, we additionally analyzed non-parametric distribution parameters, namely skewness and kurtosis, between the groups. For both parameters and studies, analysis revealed no differences in these distribution parameters (3T: all ANOVA F(4,14)<1.04, p>.4; 7T: all ANOVA F(4,9)<1.2, p>.4, all Cohen’s d<.39).

The mean values of the MDF values do vary according to the intervention groups with higher motion in all NF groups.

#### 3.1.2 Condition-wise comparisons (Does motion vary as a function of regulation condition?)

For both studies and within each group, we tested whether the MDF values vary according to the regulation conditions. ANOVA testing revealed differences in head motion between HAPPY, COUNT and REST. The post-hoc t-tests have shown in all groups increased head motion during the REST condition. For readability, we summarized all statistical values in Table 1 presenting the mean values, standard deviations for each group and condition combined with ANOVA F- and p-value. Additionally, the directionality of the differences and their effect size are summarized in the last column. The MDF values vary as function of the regulation condition in all groups of both studies regardless of the NF intervention.

**Table 1:** ANOVA statistics with F and p values and post-hoc t-test results showing directions of differences in head motion within the groups. All our groups revealed significant differences between the regulation conditions while head motion is higher during REST blocks in all groups (condition with increased mean always named first).

| Condition/<br>group | HAPPY |  | COUNT |  | REST |  | F(2,8) | p= | post-hoc p & Cohen's d |
| --- | --- | --- | --- | --- | --- | --- | --- | --- | --- |
|  | mean | std | mean | std | mean | std |  |  |  |
| 7T INT | .085 | .002 | .078 | .004 | .083 | .007 | 6.93 | .02 | HAPPY vs COUNT p=.002, d=2.2<br>REST vs. COUNT p=.004, d=.87 |
| 7T NOF | .057 | .003 | .057 | .004 | .063 | .003 | 17.4 | .001 | REST vs. HAPPY p=.002, d=2<br>REST vs. COUNT p=.004, d=1.7 |
| 3T INT | .076 | .004 | .083 | .01 | .092 | .004 | 15.2 | .002 | REST vs. HAPPY p=.002, d=4 |
| 3T NOF | .054 | .007 | .059 | .007 | .067 | .013 | 9.46 | .008 | REST vs. HAPPY p=.007, d=1.2 |
| 3T CON | .078 | .001 | .077 | .007 | .087 | .008 | 12.3 | .004 | REST vs. HAPPY p=.01, d=0.99<br>REST vs. COUNT p=.006, d=1.3 |

#### 3.1.3 Dynamics of motion (Does motion vary over time?)

To test whether within the subjects the head motion is changing over time, we tested within each group for individual differences along the five runs. Notably, for all groups there are no changes in MDF values within the groups along time (ANOVAs 7T INT F(4,95)=.12, 7T NOF F(4,65)=.16, 3T INT F(4,85)=.49, 3T NOF F(4,35)=.57, 3T CON F(4,75)=.87; all p>.48, all Cohen’s d < .31). The changes along the NF training runs are illustrated for each participant along the rows of Figure 1 in each group. We found no significant changes in head motion over time within participants.

#### 3.1.4 Correlation Regulation capability and head motion (Does motion correlate with learning effects?)

We calculated the Spearman correlations between the up-regulation success of the amygdala BOLD signal with the median framewise displacement values according to the groups collapsed across both studies (INT Baseline: r_s_=-.24, p=.15; run1: r_s_=-.1, p=.54; run2: r_s_=-.37, p=.02; run3: r_s_=-.19, p=.26; transfer: r_s_=.13, p=.46; NOF Baseline: r_s_=-.24, p=.3; run1: r_s_=-.17, p=.45; run2: r_s_=-.21, p=.35; run3: r_s_=-.27, p=.25; transfer: r_s_=-.15, p=.51). Taking Bonferroni corrections for multiple comparisons into account, Spearman correlations between the up-regulation success and the amount of head motion did not reach significance.

### 3.2 Physiological data differences (How do physiological parameters vary during the neurofeedback experiment?)

Comparable to the median framewise displacements, we analyzed differences in the physiological parameters of HR, HRV and RVT using ANOVA testing within and between groups.

#### 3.2.1 Overall group comparisons (Do they vary as a function of intervention group?)

Our ANOVA analysis comparing physiological parameters between groups revealed a significantly higher heart rate for the NOF group only with a small effect size (HR: mean INT=.87±.12, mean NOF=.94±.17, F(1,4)=363, p<.001, Cohen’s d=.49). HRV and RVT did not show differences between the groups (HRV mean INT=.0627±.03, mean NOF=.0632±.032, F(1,4)=.02, p=.86, Cohen’s d=.02; RVT mean INT=.44±.15, mean NOF=.48±.16, F(1,4)=5.9, p=.07, Cohen’s d=.26). The distribution of all parameters within both groups are shown as violin plots in Figure 4 with HR (Fig 4A), HRV (Fig. 4B) and RVT (Fig. 4C), each compared to HCP data (see 3.3)

#### 3.2.2 Condition-wise comparisons (Do they vary as a function of regulation condition?)

Our analysis revealed differences in HR, HRV and RVT between the conditions HAPPY, COUNT and REST periods in both INT and NOF group independently. Table 2 summarizes all average values and standard deviation together with ANOVA statistics and post-hoc t-tests. RVT values show lower values during COUNT condition in both groups, while HR is increased during COUNT compared to both other conditions. Further

**Table 2.** ANOVA statistics with F and p values and post-hoc t-test results showing directions of differences in HR, HRV and RVT within the groups. All our groups revealed significant differences between the regulation conditions with inconsistent directions of differences and small effect sizes.

| Condition/<br>group | HAPPY |  | COUNT |  | REST |  |  |  |  |
| --- | --- | --- | --- | --- | --- | --- | --- | --- | --- |
| Heart Rate (HR) |  |  |  |  |  |  |  |  |  |
|  | mean | std | mean | Std | mean | std | F(2,8) | p= | post-hoc p & Cohen's d |
| 7T INT | .88 | .13 | .87 | .11 | .86 | .12 | 5.5 | .03 | HAPPY vs REST p=.04, d=.17 |
| 7T NOF | .93 | .16 | .95 | .16 | .94 | .17 | 46.4 | .001 | COUNT vs. HAPPY p<.001, d=.12<br>COUNT vs. REST p<.001, d=.06 |
| Heart rate variability (HRV) |  |  |  |  |  |  |  |  |  |
| 7T INT | .06 | .028 | .067 | .031 | .061 | .032 | 7.53 | .015 | COUNT vs HAPPY p=.02, d= |
| 7T NOF | .06 | .031 | .065 | .033 | .064 | .033 | 14.92 | .002 | REST vs. HAPPY p=.01, d=<br>HAPPY vs. COUNT p=.003, d= |
| Respiratory volume per time unit (RVT) |  |  |  |  |  |  |  |  |  |
| 7T INT | .44 | .15 | .43 | .15 | .46 | .16 | 15.2 | .002 | HAPPY vs COUNT p=.05, d=<br>REST vs. COUNT p=.002, d=. |
| 7T NOF | .48 | .16 | .45 | .15 | .5 | .16 | 44.5 | <.001 | HAPPY vs. COUNT p<.001, d=<br>REST vs. COUNT p<.001, d= |

**Table 3:**
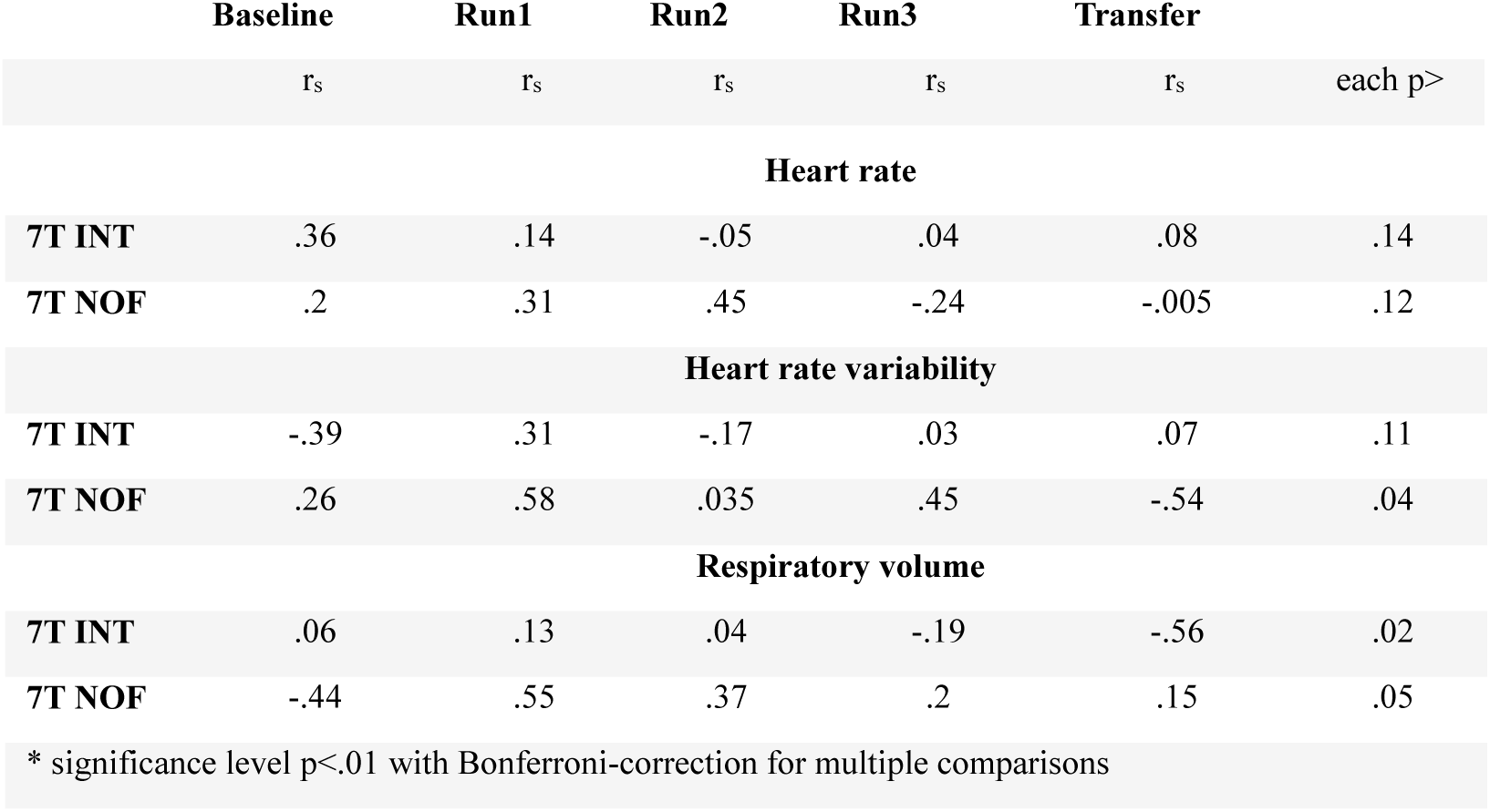
Correlation values between amygdala extracted regulation success in BOLD % signal changes and physiological noise variables along all runs.

#### 3.2.3 Dynamics of motion (Do they vary over time?)

To test whether within the subjects HR, HRV or RVT are changing over time, we tested within each group for individual differences along the five runs. Notably, for all groups and parameters we found no changes within the groups along time (all 7T INT F(4,93)<.87, p>.48, 7T NOF all F(4,65)<.99, p>.47, all Cohen’s d <.3). The physiological parameters do not vary over time.

#### 3.2.4 Correlation Regulation capability and head motion (Do they correlate with learning effects?)

We calculated the Spearman correlations between the up-regulation capability with HR, HRV and respiration values (see summary in Table 2) for INT and NOF group along all runs.

Comparable to the median framewise displacements, we revealed no correlations between amygdala regulation success from BOLD % signal with any of the physiological parameters that might explain the NF training effects in the INT group.

### 3.3 Validation analysis HCP data

In order to assess whether our findings for MFD and physiological parameters fall within standard parameter variability we examined data from HCP. Furthermore, we used these data to examine whether parameter changes revealed from our data correspond to parameter changes observed in different tasks in HCP data, where these tasks differed in levels of required mental engagement. Therefore, we used a big sample of HCP data comprising REST1, EMOTION, and MOTOR tasks. We extracted and analyzed MFD values and physiological parameters from the HCP sample data for each task analogue to our data sets. Overall, all parameter values we obtained for our small sample size are within the distribution of parameter ranges observed from the big HCP dataset. With regard to MFD values, we found significantly higher MFD values in MOTOR vs. REST1 (difference in MFD=.007 mm, p<.001) and EMOTION vs. REST1 (difference in MFD=.011, p<.001). The higher head motion between EMOTION and REST1 data from HCP mirrors our findings between feedback groups (INT, CON) and control groups (NOF), although the NOF groups performed the mental tasks of happy memories and counting backwards but without any visual feedback. Figure 3 summarizes the MFD values for all three samples (7T, 3T and HCP) averaged across all runs within each participant (Note that MOTOR task is not illustrated since results are similar to EMOTION).

**Fig. 2.**
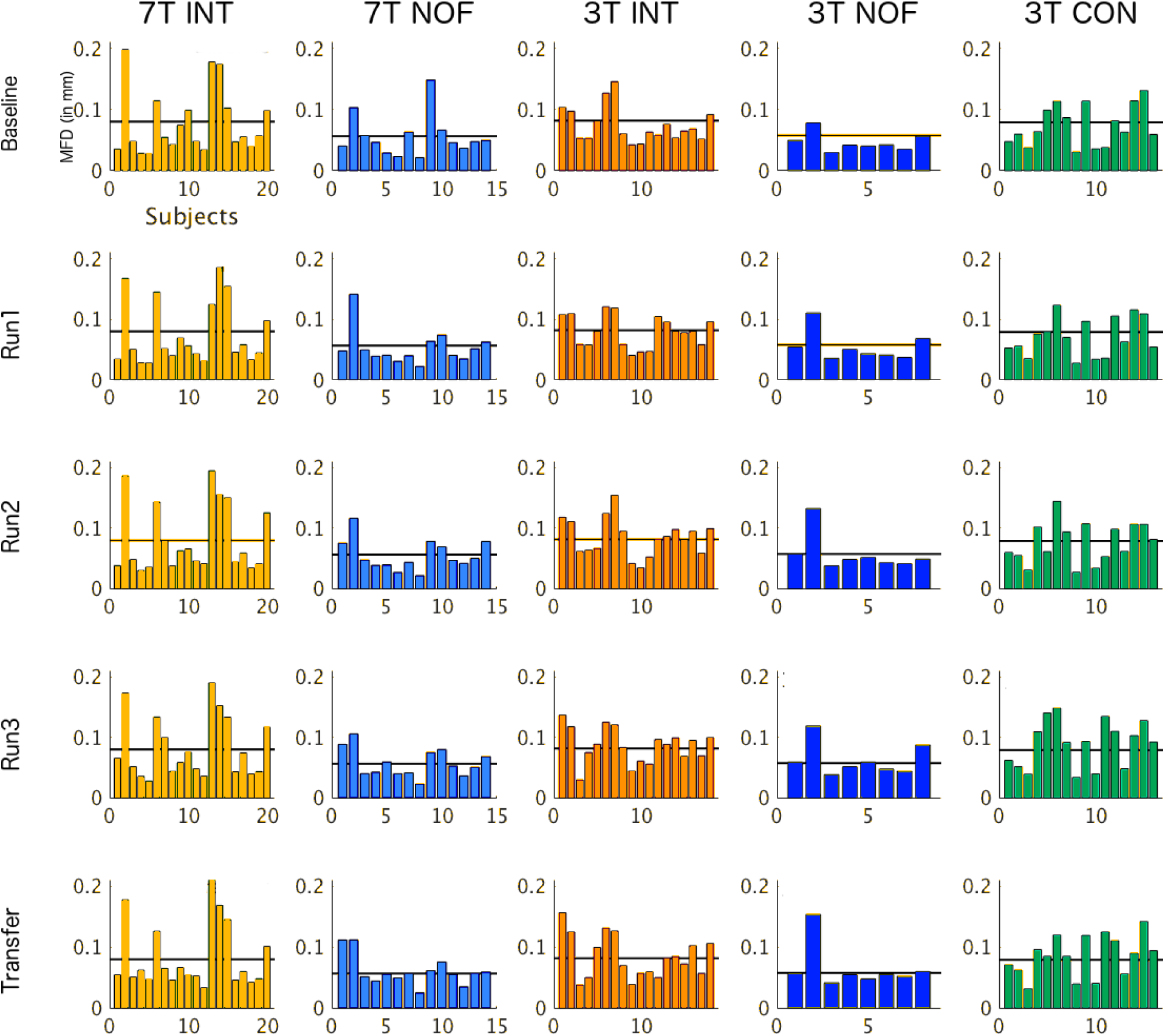
Median frame-wise displacement (MFD) in mm is plotted for every subject for (A) 7T intermittent feedback, (B) 7T no feedback, (C) 3T intermittent feedback, (D) 3T no feedback, and (E) 3T control group. The differences along the runs of the NF paradigm are plotted along the rows. The black horizontal line indicates the group average of MFD values within the respective group.

**Fig. 3.**
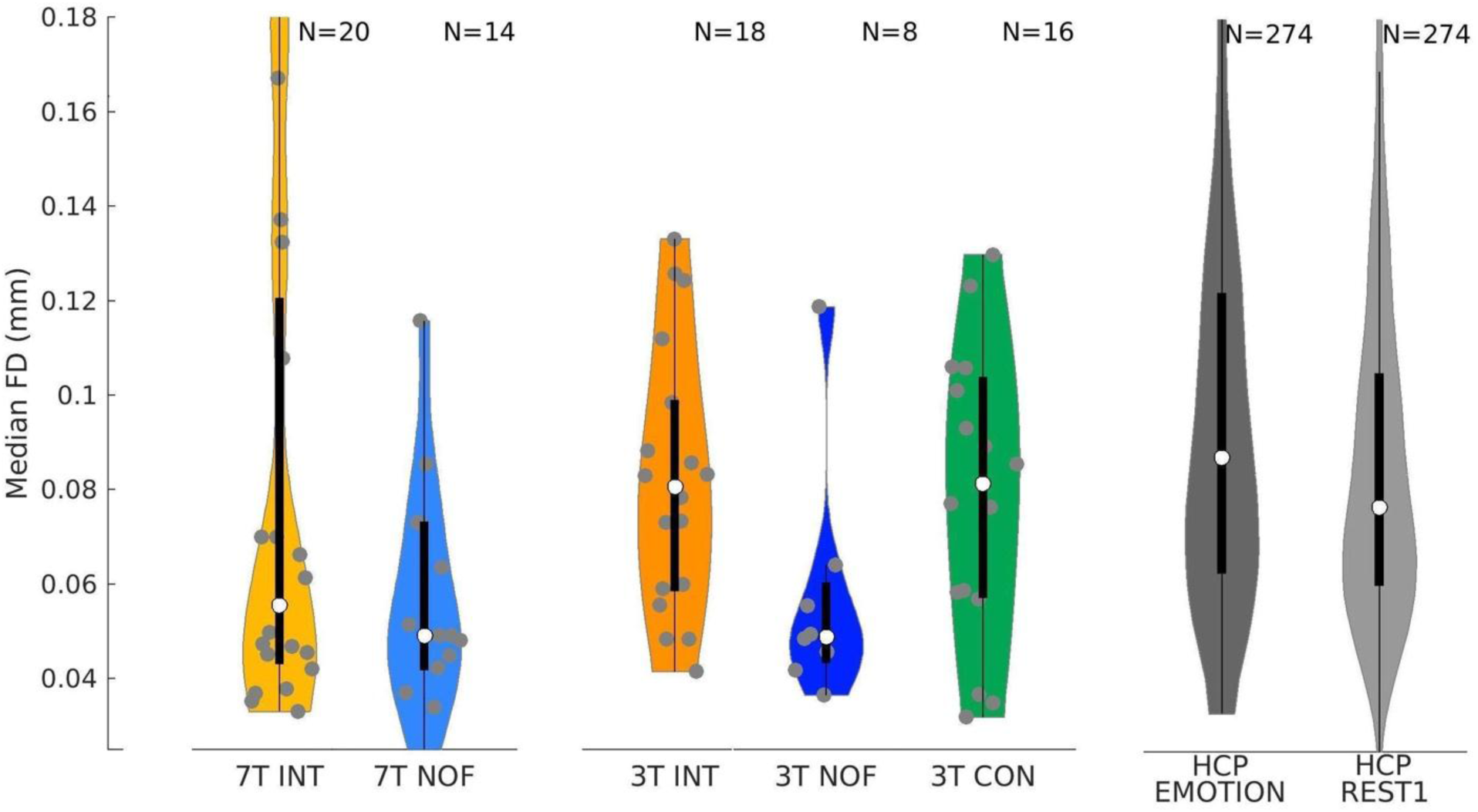
Comparison of movement in our data with the HCP data. The MFD parameter ranges we observed in our data sets fall within the normal range of parameter values. Moreover, task-based fMRI resulted in higher MFD values than resting-state fMRI in the HCP data. This finding validates our findings from both rt-fMRI amygdala NF data sets with a bigger sample size.

With regard to physiological parameters, HCP data revealed significantly higher HR values in both EMOTION and MOTOR compared to REST1 (difference EMOTION vs. REST1 = 2.2 bpm, difference MOTOR vs. REST1 = 1.5 bpm, p<.001). Furthermore, we found lower HRV in EMOTION compared to REST (difference=-.017) but higher HRV in MOTOR vs. REST (difference=.007, p<.001). For RVT, both tasks revealed significantly higher values in both tasks compared to REST1 (difference=.16; =.07; p<.001). Figure 4 summarizes heart rate (A), HRV (B) and RVT (C) values from our 7T dataset and HCP sample averaged across all runs within each subject.

**Fig. 4.**
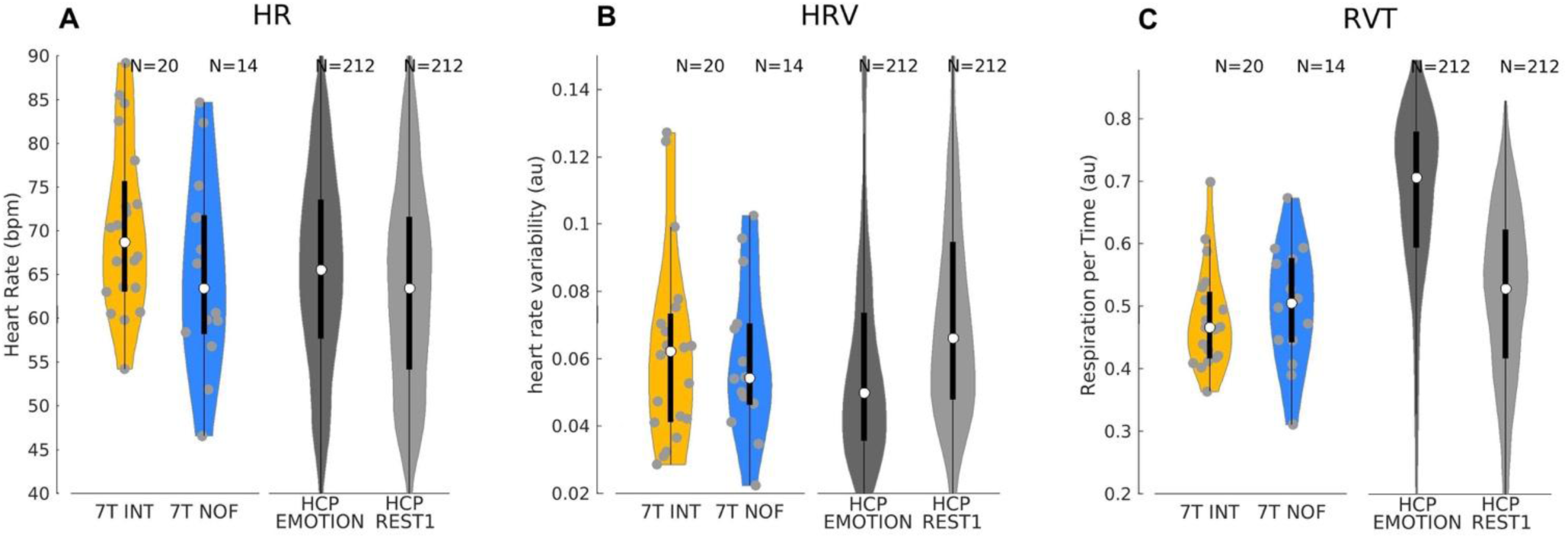
Comparison of physiological parameters in our data with the HCP data. To validate our findings for heart rate, heart rate variability and respiration volume, we analyzed the emotional task and resting state data from the HCP project. The parameter ranges in our data were comparable to those of the HCP sample. Second, although our sample did reveal significant differences in HR only with a small effect size, the bigger sample from the HCP data revealed significant differences in heart rate (A), heart rate variability (B) and respiration volume (C).

### 3.4 Amygdala task–motion and task–physiological interactions (How does the main outcome of an rt-fMRI study potentially change when motion and physiological parameters have been accounted for?)

#### 3.4.1 Exemplary single participant comparison

Exemplarily, we show a representative participant from the 7T INT group comparing amygdala % signal change extractions from the three GLMs including different numbers of nuisance regressors from no nuisance regressors over motion regressors only to motion combined with physiological regressors – GLM NO NUISANCE (Fig. 5 left), GLM + MOCO (Fig. 5 middle), and GLM + MOCO & PHYSIO (Fig. 5 right). For this participant, the comparison between GLM NO NUISANCE and GLM + MOCO & PHYSIO revealed no differences between the extracted values (p=.46). We inspected these values from all our participants’ data observing changes in the absolute values between the different GLMs. Therefore, we collapsed these values across INT and NOF groups to answer our question whether estimated group learning effects will be affected by approaches to correct for motion and physiological confounds.

**Fig. 5.**
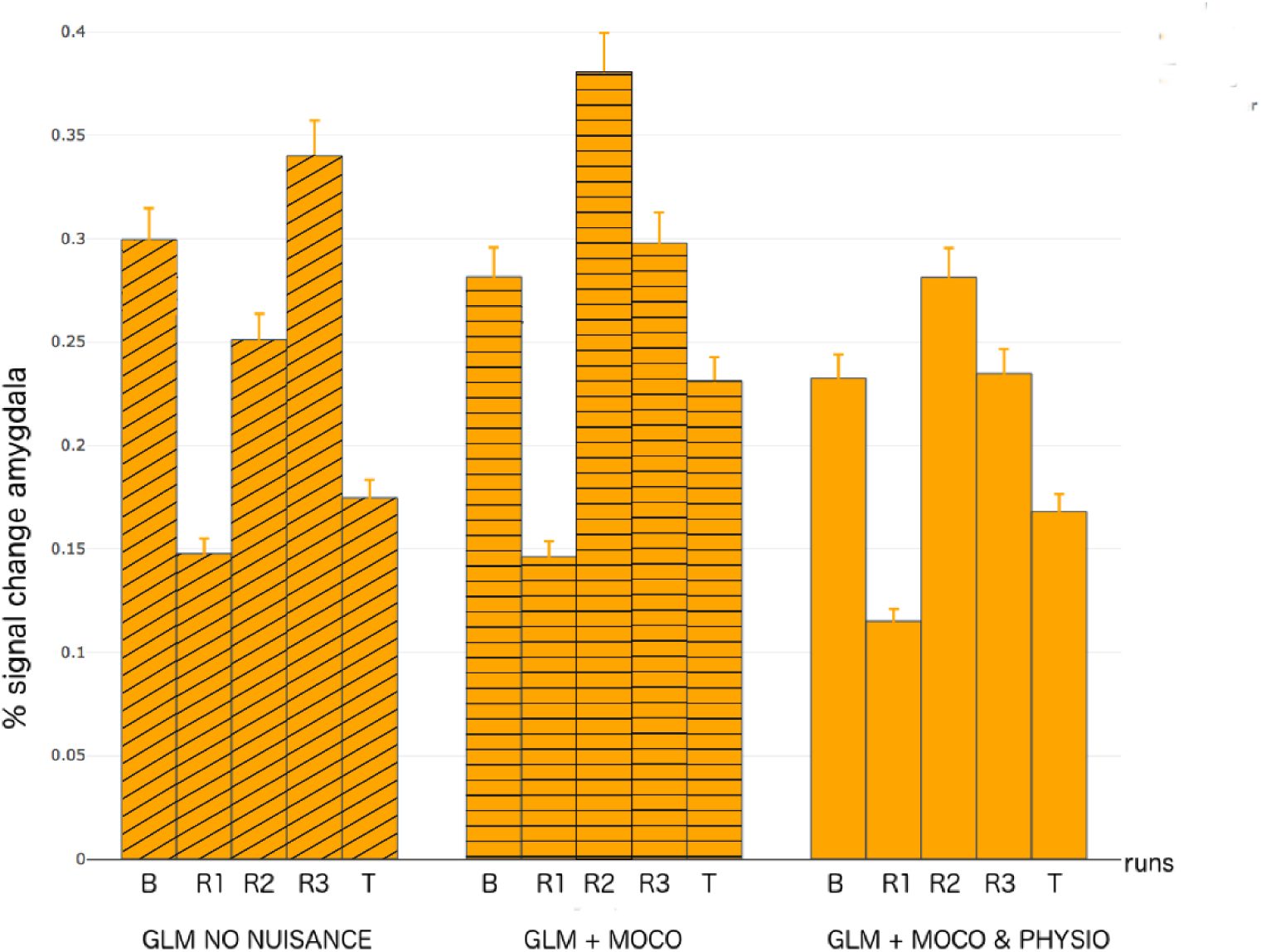
Single participant comparison of learning effects: Comparison of GLMs comprising no nuisance regression (GLM NO NUISANCE), motion regression only (GLM MOCO), and motion and physiological noise regression (GLM MOCO & PHYSIO) for one participant of 7T INT group. We found no differences regarding the percentage signal change extraction values between the GLM NO NUISANCE and GLM + MOCO & PHYSIO.

#### 3.4.2 Group level comparison 7T and 3T

For group-level comparisons, we compared the % signal change in amygdala along the runs between the three different GLMs – GLM NO NUISANCE, GLM + MOCO, and GLM + MOCO & PHYSIO – using linear mixed effect model with GLM type and run as within-subject factors. Our results revealed no main effect on GLM-type for the 7T INT group (*χ*^2^ (2)=.48, p=.78). For the 7T NOF group we found a main effect on GLM-Type (*χ*^2^(2)=7.7, p=.02). For both 3T groups we found no main effects on GLM-type (*χ*^2^(1)=3.1, p=.08; *χ*^2^(1)=2.8, p=.1). Figures 6, 7 and 8 illustrate the distributions of values within each group with boxplots for the three (7T) or two (3T) GLM types along the five runs. The cyan diamonds indicate the average values. We found

**Fig. 6.**
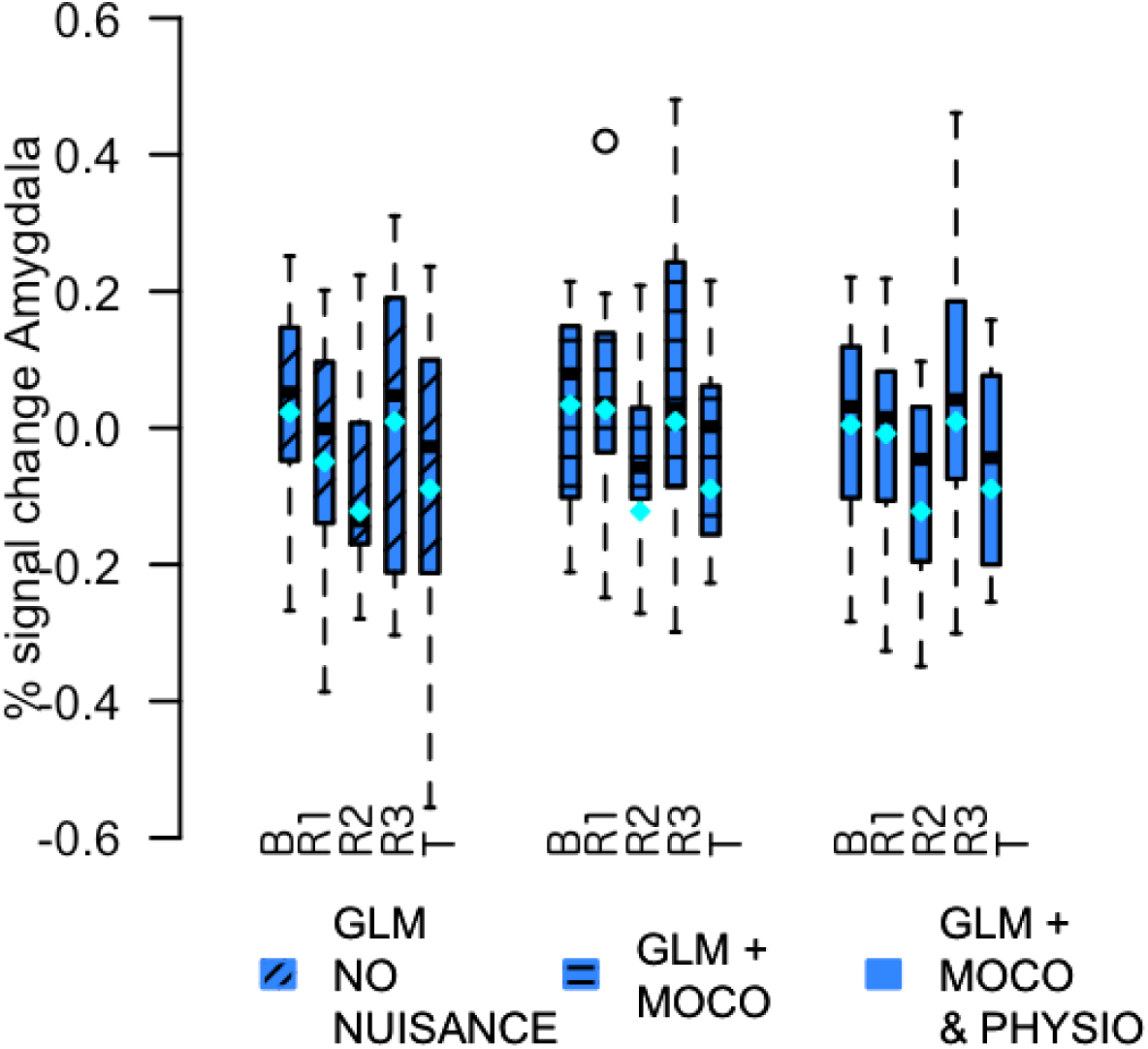
7T NOF group comparison: Comparison of amygdala extracted % signal change from GLMs without nuisance regressors, motion regressors only and motion plus physiological noise regressors. The additional regression of noise reveals a main effect of GLM comparison within this control group.

**Fig. 7.**
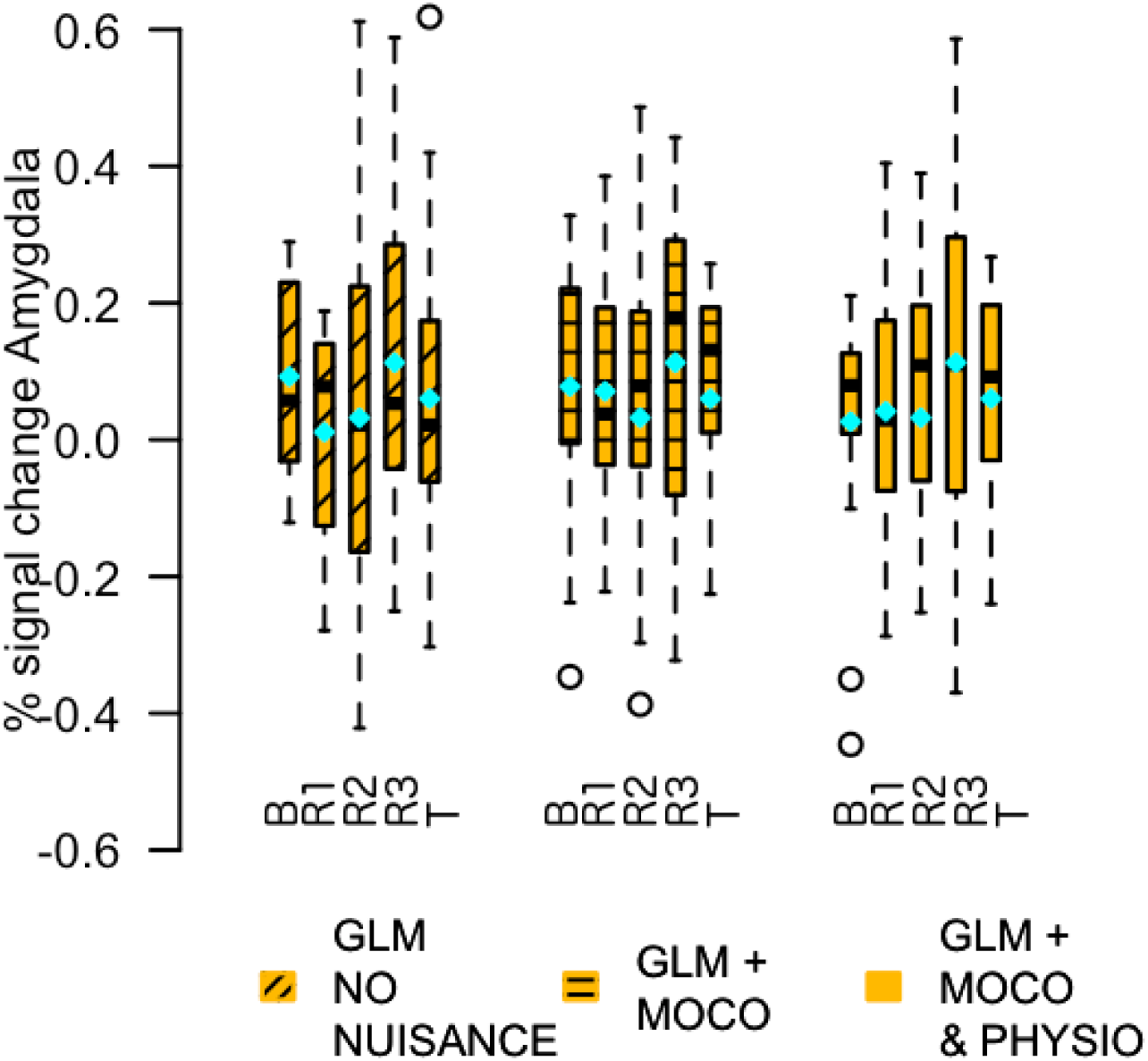
7T INT group comparison: Comparison of amygdala extracted % signal change from GLMs without nuisance regressors, motion regressors only and motion plus physiological noise regressors. The additional noise regressors did not change the results significantly.

**Fig. 8.**
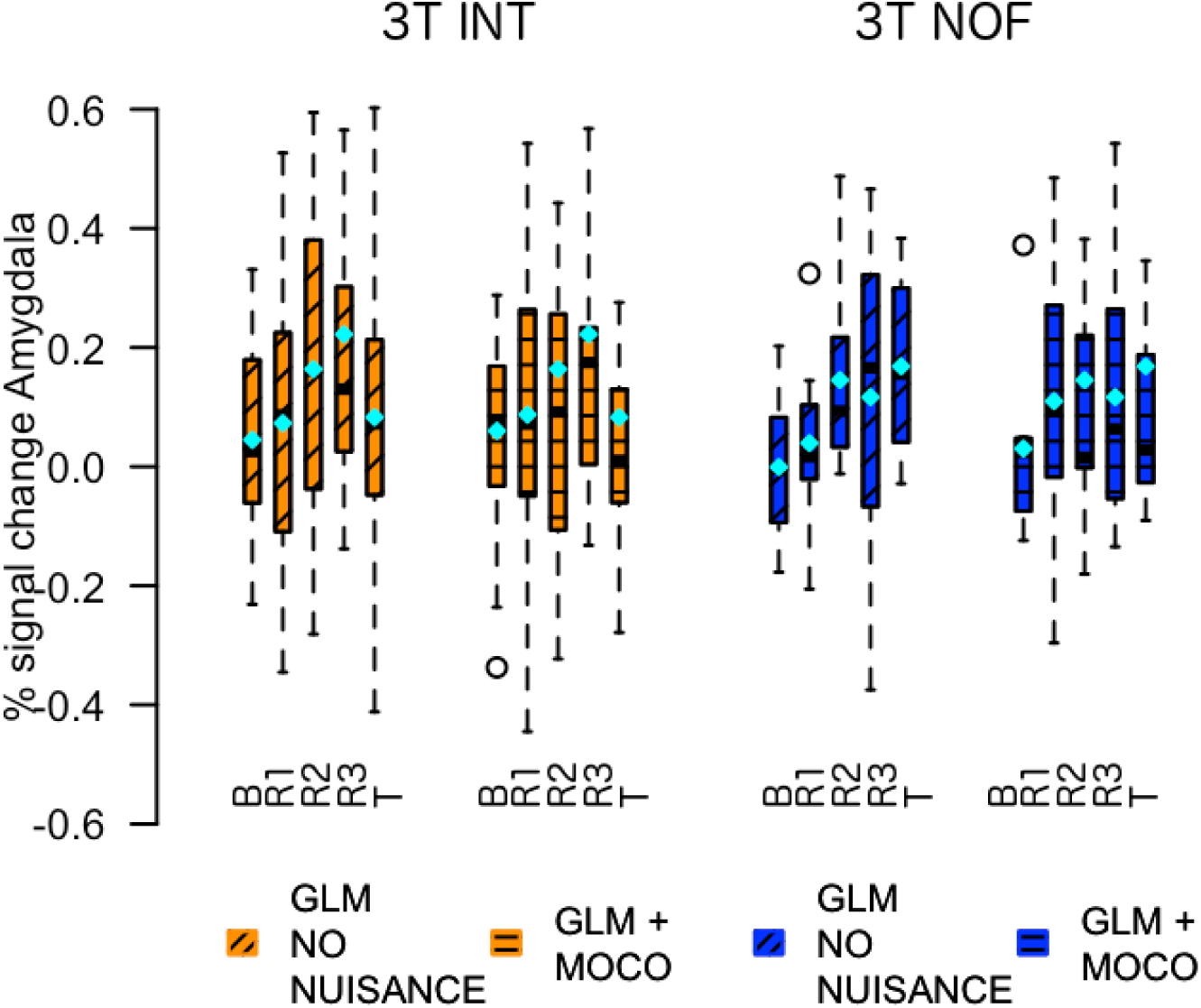
3T INT and NOF group comparison: Comparison of amygdala extracted % signal change from GLMs without nuisance regressors and motion regressors. The additional regression of motion did not change the results significantly.

#### 3.4.3 Effects of GLM enhancement on NF group differences

To answer how far statistical group differences in NF learning will be affected by such an enhanced noise correction, we performed mixed effect model analysis using INT and NOF group as between-subject factor and run as within-subject-factor. Critically, we found a difference in the resulting overall learning effects between the INT and NOF group for the comparison of amygdala signal changes. The learning effects based on the GLM NO NUISANCE would lead to significant main effects of group (*χ*^2^(1)=3.9, p=.047) and run (*χ*^2^(3)=9.6, p=.021) but no interaction. Repeating this test with the GLM + MOCO & PHYSIO evolves strong trends towards these results (main effect for group *χ*^2^(1)=2.8, p=.09 and run *χ*^2^ (3)=4.3, p=.22) and also no interaction. Our results show that the additional noise regressors on first level decreases the statistical significance of the results although the interpretation remains.

## 4 Discussion

In two independent amygdala neurofeedback studies, we systematically investigated effects of head motion, heart rate, heart rate variability and respiratory volume on BOLD signal changes achieved from neurofeedback training. We mainly asked three questions: (1) How do these parameters vary during a neurofeedback experiment and if they vary as a function of either intervention (feedback or no feedback), or regulation condition (HAPPY, COUNT, REST), or along training runs (Baseline, Run1, Run2, Run3, Transfer)? In response to this, our results revealed that motion differed between groups and between regulation condition. For physiological parameters, we found a slightly higher heart rate in the no feedback group with a small effect size. Furthermore, the parameters differ between regulation conditions inconsistently with very small effect sizes. All nuisance parameters did not vary along training runs. (2) Do these parameters correlate with learning effects, i.e. increase together with increased capacity of BOLD regulation across several training runs? Here, our results have shown that all nuisance parameters were uncorrelated to success in volitional self-regulation capability of the amygdala (see 3.1.4 and 3.2.4). (3) How does the main outcome of an rt-fMRI study (i.e. the demonstration of BOLD regulation and its transfer) potentially change when motion and physiological parameters have been accounted for? In response to this question, we found that including all nuisance regressors properly did not change the interpretation of the results but affected the statistical results in terms of a decrease in statistical power (see 3.4.3). Therefore, we conclude that overall changes in BOLD activity achieved from amygdala neurofeedback training are not mainly driven by noise. Our results underline that neurofeedback allows to learn self-regulation of brain activity reliably with ongoing training.

Our analysis of the HCP data showed that as engagement with the task or the presented stimuli increases, by going from a resting state (i.e. REST1) task to a more active task (i.e. EMOTION or MOTOR), motion and physiological parameters change significantly. In particular, motion increases, heart rate and respiration increase, whereas heart rate variability decreases. This is in line with reported literature describing the effects of these parameters on BOLD (Lund et al., 2005; Sakaki et al., 2016; Thayer, 2018). Furthermore, it has been shown that higher heart rate variability in self-control is associated with altered brain activity in ventromedial prefrontal cortex but not amygdala (Maier and Hare, 2017). The results in our studies confirm to these findings, especially as we found in both groups only within-subject differences between the regulation conditions.

The combined evidence indicates that the reason behind the observed differences between our groups might be the feedback information, which might lead to a stronger engagement of the participants in the task. The presence of the feedback and its interpretation adds a cognitive process for the participants, since the level of the stimulus needs to be evaluated/consolidated with the own current belief of their performance. Furthermore, this allows to evaluate whether the strategy employed before the feedback aroused needs to be changed. This entails the process of learning over BOLD signal even with different kinds of feedback – either presented continuously or occasionally only every 40 seconds. Critically to our study design, we used only one control group instructed identical with regard to strategies but receiving no feedback at all. But with regard to behavioral differences, we found no differences in strategy usage or willingness to perform the task between our groups. The compliance of the control groups is also underpinned by our findings from HCP analysis revealing significant differences between resting-state and task-fMRI, while our groups do not differ significantly in physiological parameters.

One could argue that the amygdala BOLD effects may be mainly driven by these physiological factors, especially considering the publication of Boubela et al. (2015) warning exactly for these possible effects. It therefore is especially essential to control for these confounders. Although in our results this is clearly not the case as the amygdala BOLD variability across participants is not correlated with any motion or physiological parameters, potential confounds of head motion and physiological parameters needs to be taken into account, especially when analyzing average extracted amygdala BOLD effects. When they have been extracted with a GLM approach that includes motion and physiology, the degrees of freedom at first level are decreased due to the additional motion (N=12) and physiological (N=8) regressors, which could lead to a decreased statistical power when assessing group differences (Faul et al., 2009). Therefore, an appropriate sample size is needed.

Regarding motion, there is a conflicting report showing decreased motion going from resting-state to task-based fMRI (Huijbers et al., 2017), which conflicts with findings from our own studies (3T and 7T) and also (more clearly) conflicts with our findings from the analysis of 822 participants from the HCP. However, Huijbers at al. state the possible effects of different kinds of cognitive tasks they used, use mainly clinical population from different psychiatric disorders and age effects on their reported motion values. The HCP data comprise a lower age range with overall less motion and comprises healthy adult participants.

Limitations: A critical aspect to our study might be that the different physiological and motion signals we assess come with different time resolutions. Due to the nature of fMRI the correction methods we have used are within the repetition time of our sequence (2s). Aside, the data we acquired are in high resolution (1.4 mm isotropic) at ultra-highfield strength, which has been shown to be more sensitive to physiological noise than standard 3T sequences, although different parts of the brain are affected differently by physiological changes (van der Zwaag et al., 2015). However, we used the current state-of-the-art methods for data acquisition and nuisance correction. Overall, our results cannot be fully generalized to all kinds of rt-fMRI neurofeedback studies and it is crucial that more fMRI-NF studies report noise corrected results.

Although our study underpins the reliability of neurofeedback allowing to increase BOLD activity over the training, we think it is highly recommended to acquire motion and physiological data, and also highlights the importance of post-hoc offline analysis taking these into account when analyzing the data. This would help to increase trust in all studies reporting neurofeedback training effects. However, our results indicate that there is not crucial alteration of the main conclusions of the experiment, and therefore, no pressing need for the rt-fMRI field to invest in the implementation of a real-time motion and physiological correction for the on-line extraction of BOLD activity that is used as the neurofeedback in the experiment. At the moment, this would be only possible if an EEG system is used that supports real-time data export (for example Brain Products Recorder or EGI Netstation). EEG systems are not always available and would increase the complexity of the setup unnecessarily. Second, a real-time analysis would have to be performed to extract the physiological parameters such heart rate and respiration rate, before they would be able to be used as regressors for on-line statistics.

Nevertheless, our recommendation for further studies is to carefully report in addition to study results a scheme of neurofeedback learning effects with post-hoc regression of physiological noise parameters to prove the validity of the changes in neural activity.

## 5 Conclusions

We conclude that motion artifacts did not drive the improvement in self-regulation capability of the amygdala and other feedback related brain regions. But the differences in motion parameters between feedback and control groups emphasize the importance to control for such confounders. Furthermore, we strongly suggest to monitor these parameters, such as head movement, pulse and respiration during real-time fMRI experiments and to report these data for strengthening the effects of rt-fMRI neurofeedback interventions. Moreover, to account for motion differences within subjects and between groups, we propose to use analyses that comprise the median frame-wise displacement values and neurofeedback effects such that physiological variability is regressed out to precisely show self-regulation improvements in neurofeedback experiments.

## Acknowledgements

The authors would like to thank Renate Blobel for all the help in recruitment and scanning at the 7T MR scanner. We also thank Maria Watermann and Maya Nathan for help with recruitment. We also thank Myung-Ho In from the Department of Biomedical Magnetic Resonance, Magdeburg, for getting started with the MR sequence. Furthermore, we thank Saskia Bollmann for personal help with the usage of TapasIO Toolbox. This work was supported by grant 00014_165884 from the Swiss National Science Foundation (to PNT) and from the European Union’s Horizon 2020 research and innovation programme under the Marie Sklodowska-Curie grant agreement 794395 (to LH).

The authors declare no competing financial interests.

## Footnotes

1 http://fsl.fmrib.ox.ac.uk/

2 https://cni.lin-magdeburg.de/index.php/en/wiki/mri/physiological-data/

3 http://www.mathworks.com

